# Virucidal activity of CPC-containing oral rinses against SARS-CoV-2 variants and are active in the presence of human saliva

**DOI:** 10.1101/2021.08.05.455040

**Authors:** Enyia R Anderson, Edward I Patterson, Siobhan Richards, Alison Green, Sayandip Mukherjee, Michael Hoptroff, Grant L Hughes

## Abstract

The role of human saliva in aerosol-based transmission of SARS-CoV-2 has highlighted the need to understand the potential of oral hygiene products to inactivate the virus. Here we examined the efficacy of mouthwashes containing cetylpyridinium chloride (CPC) or chlorhexidine (CHX) in inactivating SARS-CoV-2. After 30 seconds contact under standard aqueous conditions CPC mouthwashes achieved a ≥4.0log_10_ PFU/mL reduction in SARS-CoV-2 (USA-WA1/2020) titres whereas comparable products containing CHX achieved <2.0log_10_ PFU/mL reduction. Further testing with CPC mouthwashes demonstrated efficacy against multiple SARS-CoV-2 variants, with inactivation below the limit of detection observed against the Alpha (B.1.1.7), Beta (B.1.351) and Gamma (P.1) variants. Virucidal efficacy of CPC mouthwash was also observed in the presence of human saliva with the product delivering ≥4.0log_10_ PFU/mL reduction in SARS-CoV-2 titres after 30 seconds providing additional evidence for the virucidal efficacy of CPC mouthwashes under simulated physiological conditions. Together these data suggest CPC-based mouthwashes are effective at inactivating SARS-CoV-2 and further supports the use of mouthwash to mitigate the risk of transmission during dentistry procedures.

## Introduction

The high viral load of SARS-CoV-2 present in the saliva of infected individuals and their ease of aerosolisation means that saliva is recognised as playing a key role in the transmission of SARS-CoV-2 [1–3]. The potential of the oral cavity to act as a viral reservoir is supported by the presence of the angiotensin-converting enzyme 2 (ACE-2) receptor in oral gingival epithelia and salivary glands and the infection of these tissues by SARS-CoV-2 *in vivo* [4], potentially aggravating systemic infection via an oral-vasculo-pulmonary route [5].

The use of oral rinses or mouthwashes have been proposed by health organisations to mitigate transmission of SARS-CoV-2 during dentistry procedures due to their demonstrated efficacy in deactivating SARS-CoV-2 *in vitro* and *in vivo* [6–8]. The antimicrobial action of a mouthwash is dependent on a combination of the active ingredients, their intrinsic efficacy, and their bioavailability during use. Active ingredients used in mouthwashes include Quaternary Ammonium Compounds (QACS) such as Dequalinium chloride, benzalkonium chloride, cetyl pyridinium chloride (CPC) and chlorhexidine (CHX) which are believed function as antimicrobials via a stepwise process of charge mediated attraction and destabilisation of the lipid envelop [9–11].

CPC is widely used in mouthwash formulations displaying substantive action against a range of oral bacteria [12–14] and viruses, including SARS-CoV-2 [6, 15–17], whilst data on antiviral efficacy of CHX against SARS-CoV-2 has been more varied [18–22]. All mouthwashes, regardless of composition, must function *in situ* in the oral cavity, and hence must retain efficacy in the presence of human saliva, overcoming any potential deactivation from salivary components [23–25].

To investigate the impact of formulation composition on efficacy we compared the *in vitro* virucidal efficacy of mouthwashes containing 0.07% CPC and 0.2% CHX digluconate against a range of SARS-CoV-2 variants. In addition, the efficacy of a representative CPC containing mouthwash was also investigated in the presence of human saliva. Our findings suggest CPC mouthwashes offer potent virucidal activity that is effective against all variants tested and which is maintained in the presence of human saliva under simulated usage conditions.

## Methods

### Cell Culture and Viruses

Vero E6 cells (C1008: African green monkey kidney cells) obtained from Public Health England, were maintained in Dulbecco’s minimal essential medium (DMEM) with 10% foetal bovine serum (FBS) and 0.05mg/ml gentamicin. Cells were maintained at 37°C and 5% CO_2_. Passage 5 of SARS-CoV-2 isolate (USA-WA1/2020) and passage 2 or 3 of Alpha (hCoV-19/England/204820464/2020), Beta (hCoV-19/South Africa/KRISP-EC-K005321/2020) and Gamma (hCoV-19/Japan/TY7-503/2021) obtained from BEI Resources [26], were cultured in Vero E6 cells maintained in DMEM with 4% FBS and 0.05mg/ml gentamicin at 37°C and 5% CO_2_. 48 hours post inoculation, virus was harvested and stored at −80°C until used.

### Preparation of Saliva

Stimulated saliva was collected at Unilever Research Port Sunlight, with ethical approval from the Unilever R&D Port Sunlight Independent Ethics Committee (GEN 022 13), collected during November 2019 from 11 donors over 2 days. The saliva was pooled and aliquoted into 250ml samples, and then stored at −20°C until sterilisation. The saliva was sterilised using gamma irradiation (Systagenix, UK, Cobolt 60 turntable, Dose rate 1.2 kGy/hr, minimum dose 32.1 kGy).

### Virus Inactivation

Mouthwash formulations (Table 1 and Figure 1) were assessed following the ASTM International Standard E1052-20 [27]. SARS-CoV-2 titre was calculated for each experiment by plaque assays with the titre consistently >4.3log_10_ PFU/mL for USA-WA1/2020 and variants. Briefly, 900 μL of mouthwash formulation was added to 100 μL of virus suspension, containing 4% FBS and incubated for 30 seconds. After the 30 second incubation, 9mL of Dey and Engley neutralising broth (DE broth) was added and 25μL of the sample was transferred into a dilution series for quantification through a standard plaque assay as previously described.

**Table 1.** Mouthwash formulations examined for SARS-CoV-2 inactivation.

| Treatment | Code |
| --- | --- |
| 0.07% CPC with flavour and mix of herbal extracts | MW-A |
| 0.07% CPC with flavour | MW-B |
| 0.2% CHX digluconate with flavour | MW-C |
| 70% ethanol in distilled water | Control |

**Figure 1.**
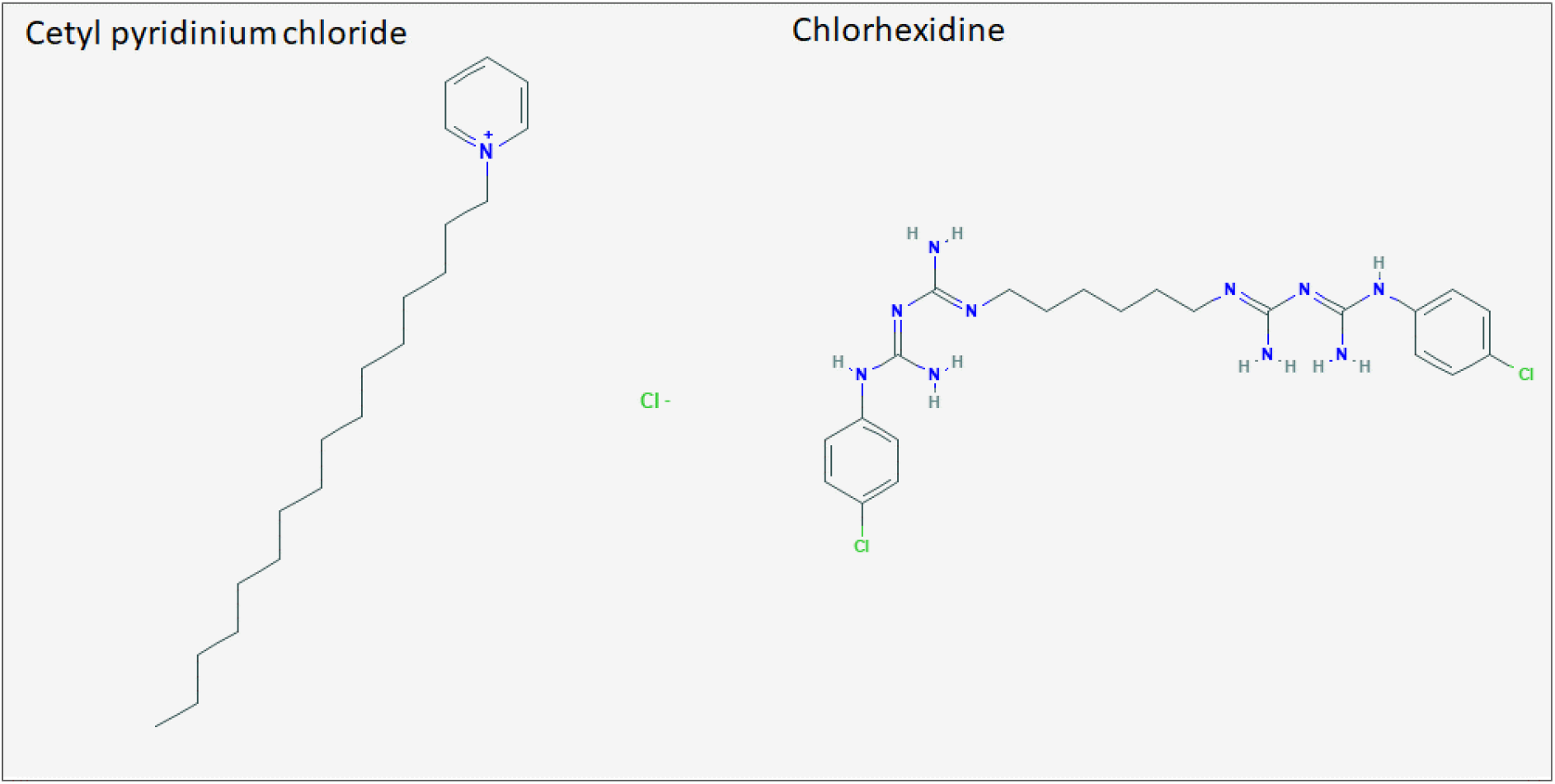
Structures of cetyl pyrinium chloride (CPC) and chlorhexidine (CHX).

To assess if human saliva alters the effectiveness of CPC mouthwash, 800 μL of MW-B mouthwash formula was added to 100 μL of human saliva mixed with 100 μL of SARS-CoV-2 (USA-WA1/2020) inoculum. The solution was incubated for 30 seconds and 9 mL of DE broth added. As a control, 800 μL of MW-B mouthwash formula was added to 100 μL of sterile water and 100 μL of virus inoculum, with 9 mL of DE broth added after 30 seconds of incubation. Experiments were carried out in duplicate.

### Saliva, neutralisation, and cytotoxicity assays

To assess if human saliva has inherent antiviral action against SARS-CoV-2 (USA-WA1/2020), 100 μL of virus inoculum was added to either 800 μL of sterile water and 100 μL of irradiated human saliva (dilute saliva), or 900μL of irradiated human saliva (neat saliva) for a 5-minute incubation. After 5 minutes had elapsed, a 25 μL sample was placed into a dilution series. Neutralisation controls were carried out by adding 9 mL of DE broth to 900 μL of mouthwash formula. To this, 100 μL of virus suspension was added for 30 seconds and 25 μL removed to a dilution series and a standard plaque assay preformed. To determine the cytotoxicity of the mouthwashes, 100 μL of 4% DMEM was added to 900 μL of test mouthwash formula for 30 seconds. To this 9 mL of DE broth was added and 25 μL placed into dilution series for a standard plaque assay. 25 μL samples from each condition were serial diluted 10-fold to be quantified through a standard plaque assay. Plaques were counted to determine viral titre. All experiments were carried out in triplicate.

## Results

### Comparison of CPC and CHX containing mouthwashes

We tested the ability of CPC and CHX to inactivate SARS-CoV-2 (USA-WA1/2020). Following a 30 second incubation in the presence of the test mouthwashes a reduction in viral titre of ≥4.0log_10_ PFU/mL was observed with MW-A and MW-B and of <2.0 log_10_ PFU/mL for MW-C. No reduction of viral titre occurred in the water control and however complete inactivation was observed by the 70% ethanol control. All treatments were effectively neutralised by the addition of DE broth (Figure 2). Cytotoxicity assays were able to determine the limit of detection (LOD) for this assay. The LOD is the point at which Vero E6 cell death is due to cytotoxicity of mouthwashes rather than SARS-CoV-2. For all three mouthwashes presented here, cytopathic effect was observed at 2.0log_10_ PFU/mL.

**Figure 2.**
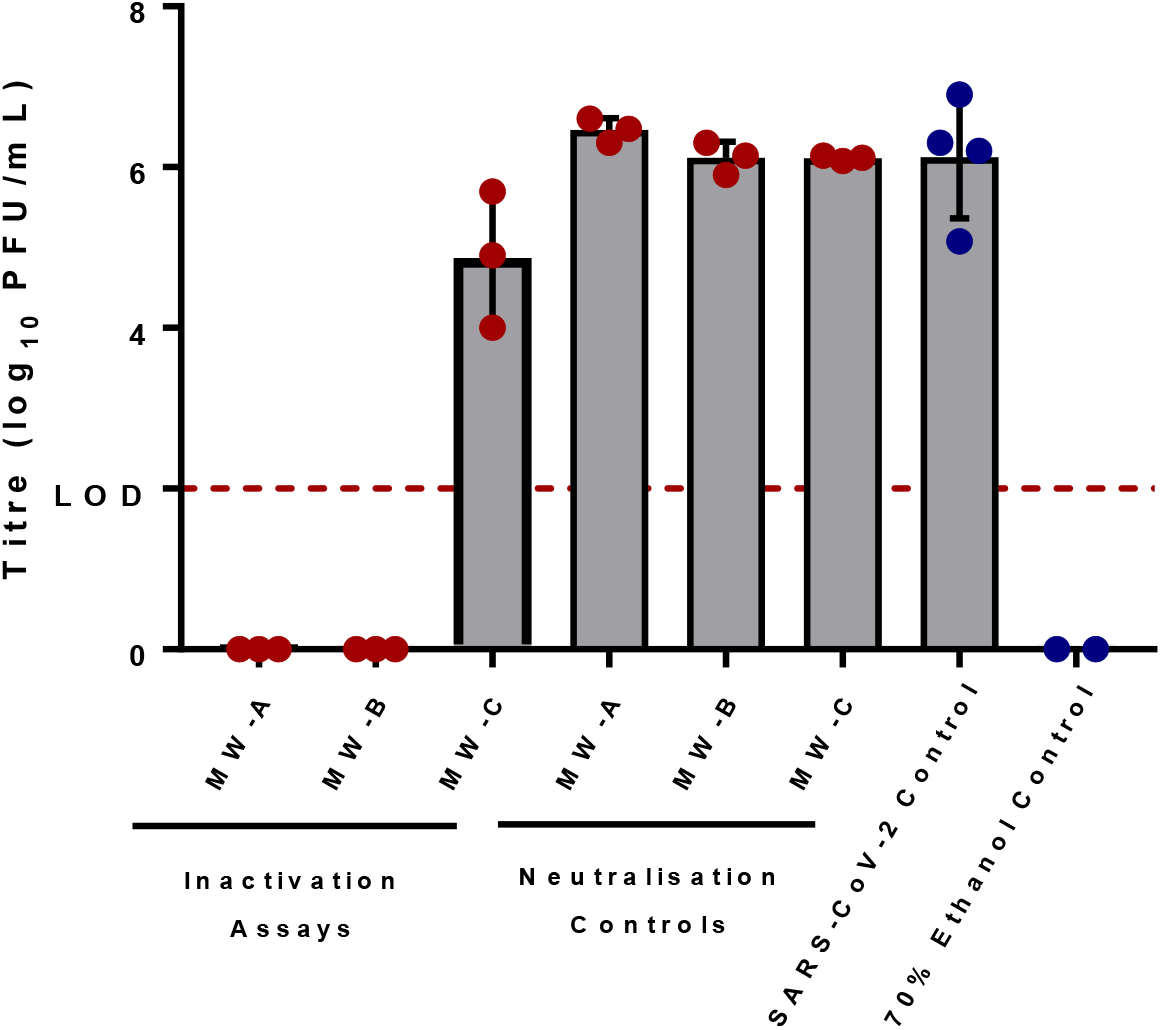
Mouthwash formulas were tested for antiviral action against SARS-CoV-2. Mouthwashes were incubated with SARS-CoV-2 inoculum for 30 seconds. Both MW-A and MW-B reduced to below the limit of detection (LOD), while MW-C reduced viral titre by 1.26log_10_ PFU/mL compared to the water control. LOD = 2.0log_10_ PFU/mL. Viral titres recovered from the water control averaged 6.12log_10_ PFU/mL, while viral titre recovered from neutralisation controls were within 1.0log_10_ PFU/mL indicating all antiviral activity occurred within 30 seconds of incubation. LOD (2.0log_10_ PFU/mL) is shown across the graph with a dotted red line. Error bars represent standard deviation, while red dots are experimental data values and blue dots control values.

### Inactivation of SARS-CoV-2 Variants by Test Products

We also tested the ability of CPC and CHX to inactivate SARS-CoV-2 variants of concern, Alpha, Beta and Gamma. Following the 30 second incubation of Alpha with MW-A and MW-B an average reduction of 3.11log_10_ PFU/mL to below the LOD was seen. Incubation of Beta with test products saw an average reduction of 4.1log_10_ PFU/mL, whilst Gamma saw an average reduction of 3.36log_10_ PFU/mL, both to below the LOD (Table 1). In assays carried out with the variants, no reduction was seen in the water control and reduction below the LOD was seen in the 70% ethanol control. The ability to achieve a 4.0log_10_ PFU/mL in the variant assays was dependent on titres of SARS-CoV-2 variants following standard propagation methods.

**Table 1.** Mouthwash formulas that were proven to work against SARS-CoV-2 (USA-WA1/2020) were then tested against Alpha, Beta, and Gamma variants of SARS-CoV-2. Both MW-A and MW-B were able to reduce the viral titre of all three variants to below the limit of detection (2.0log_10_ PFU/mL) within 30 seconds.

| Variant | Pango lineage | Average Titre (Log<sub>10</sub> PFU/mL) Reduction |  |  |
| --- | --- | --- | --- | --- |
|  |  | MW-A | MW-B | 70% Ethanol |
| Alpha | B.1.1.7 | 3.11 | 3.11 | 3.11 |
| Beta | B.1.351 | 4.11 | 4.11 | 4.11 |
| Gamma | P.1 | 3.36 | 3.36 | 3.36 |

### Testing in presence of human saliva

Under normal usage mouthwashes must be functional in the presence of human saliva, hence investigations were undertaken to assess whether saliva displays any measurable endogenous antiviral activity against SARS-CoV-2 (USA-WA1/2020) or whether it acts as an inhibitory “soil” quenching the antiviral function of mouthwash formulations. The endogenous antiviral activity of neat and dilute human saliva was measured over a contact time of 5 minutes (Figure 3A) during which no significant reduction in viral load was observed compared to the water control. Viral titres of 5.70log_10_ PFU/mL, 5.61log_10_ PFU/mL and 5.45log_10_ PFU/mL were recovered from the neat saliva, dilute saliva incubation and the water control, respectively. It is essential that mouthwashes maintain efficacy in the presence of human saliva. To investigate this, we examined if the antiviral efficacy of MW-B was altered by saliva. We found that MW-B was still capable of inactivation of SARS-CoV-2 to below the LOD in the presence of saliva, indicating that CPC retained efficacy despite the soil load.

**Figure 3.**
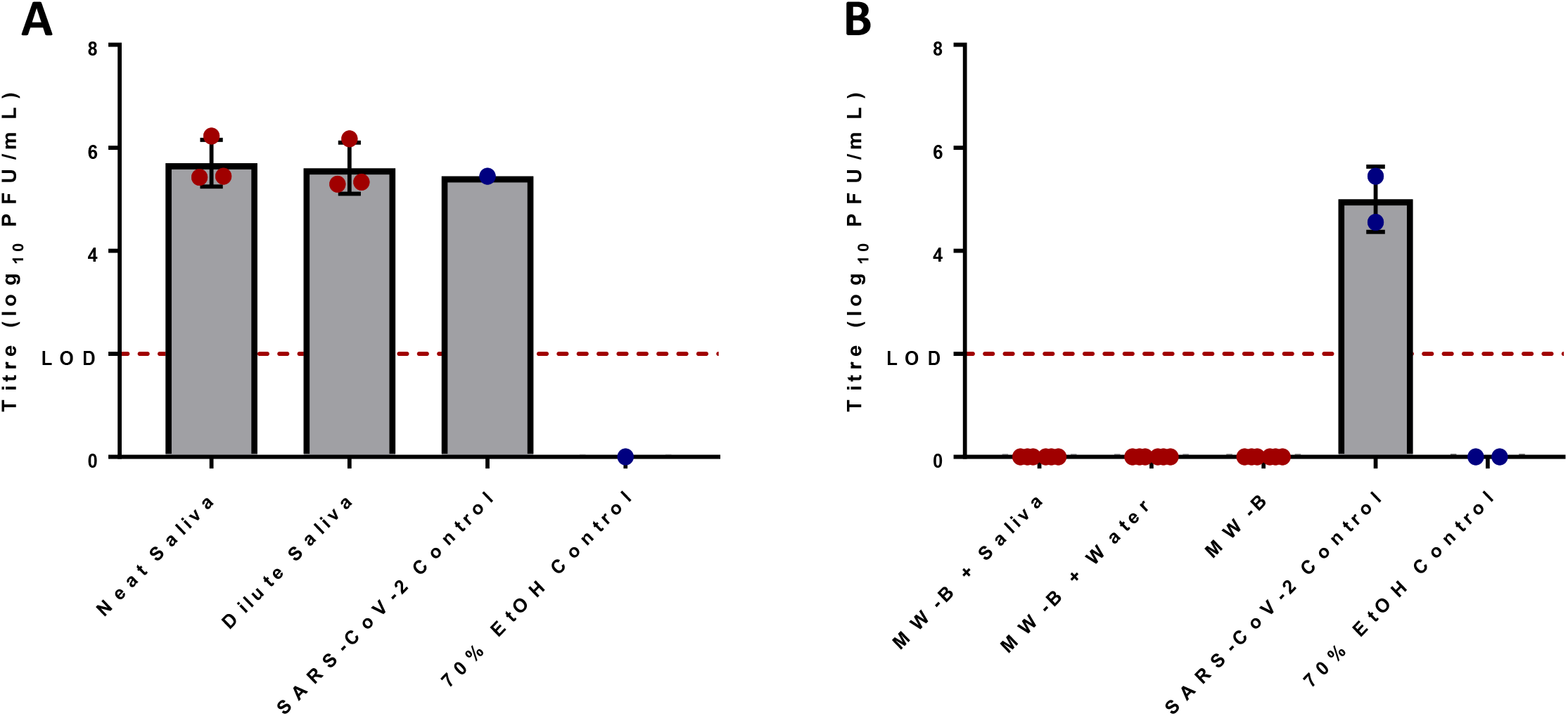
Irradiated human saliva has no effect upon the viral titre of SARS-CoV-2 as compared to the water control after incubation with inoculum for 5 minutes. Neat saliva had a ratio of 8 parts water to 1-part irradiated human saliva to 1-part virus inoculum, while dilute saliva had a ratio 9 parts irradiated human saliva to 1-part virus inoculum (A). Human saliva does not inhibit the antiviral activity of mouthwash formulas proven to reduce the titre of SARS-CoV-2 (B). MW-B was able to reduce viral titre to below the LOD both in the presence of irradiated human saliva and without. Human saliva was added in a ratio of 8 parts MW-B to 1-part irradiated human saliva to 1-part virus inoculum. Limit of detection (LOD) (2.0log_10_ PFU/mL) is shown across both graphs with a dotted red line. Error bars represent standard deviation, while red dots are experimental data values and blue dots control values.

## Discussion

Our results confirm that mouthwash formulations containing 0.07% CPC, inactivate SARS-CoV-2 by up to 99.99%, representing a value below the LOD after a contact time of 30 seconds. In contrast, within the same experiment, a mouthwash containing CHX (0.2% chlorhexidine gluconate), exhibited poorer virucidal activity against SARS-CoV-2. Our observations are consistent with others, where a number of different CPC mouthwash formulations have been shown to effectively inactivate SARS-CoV-2 *in vitro*, whereas CHX containing mouthwashes are reported to have modest ability to inactive SARS-CoV-2 [6, 19, 21]. The virucidal action of CPC mouthwash was maintained in the presence of whole human saliva, consistent with human clinical trials which report that rinsing with CPC mouthwash can lower SARS-CoV-2 salivary count for several hours after use [7, 28].

Over the course of the global pandemic, several SARS-CoV-2 variants have emerged with mutations changing the amino acid sequence of the receptor-binding domain of the spike protein [29]. Three variants of concern; Alpha, Beta, and Gamma [30] were effectively inactivated within 30 seconds by both 0.07% CPC mouthwashes, with a reduction in viral titre below the LOD and equivalent to the 70% ethanol control. As the CPC molecule disrupts the viral lipid envelope and the membrane is unchanged by mutations, our data supports the likely efficacy of CPC mouthwash in reducing viral load irrespective of the SARS-CoV-2 variant. Recently the oral cavity has been proposed to have a direct role in COVID-19 disease severity based on a proposed oral-vasculo-pulmonary infection route. Poor oral hygiene with plaque build-up, subsequent gingivitis and periodontitis facilitates direct entry of the virus via the oral gingival sulcus and periodontal pockets enabling infection of the circulatory system and lungs [5]. CPC mouthwashes with anti-plaque and virucidal activity against SARS-CoV-2 could have the potential to lower viral count and lessen the risk of severe lung disease in COVID-19 patients.

In conclusion, two mouthwashes containing 0.07% CPC were effective at inactivating SARS-CoV-2, within 30 seconds with greater than 4.0log_10_ PFU/ml reduction in viral titre. Moreover, virucidal activity of CPC was maintained in the presence of whole human saliva. Both 0.07% CPC mouthwashes were as effective as 70% ethanol against three variants of concern; Alpha, Beta and Gamma suggesting these CPC formulations possess virucidal action against all variants. In contrast, under the same experimental conditions, a mouthwash containing 0.2% chlorhexidine digluconate did not have substantial action against SARS-CoV-2 *in vitro*. Given the ongoing global pandemic, and the recognition of the significance of the oral cavity in infection, transmission, and disease severity, daily use of an effective CPC mouthwash as part of a good oral hygiene routine, could be a low-cost and simple measure to reduce transmission risk and potentially, lower the risk of developing severe forms of COVID-19.

## Acknowledgements

Unilever funded this study. EIP and GLH were supported by the EPSRC (V043811/1) and UKRI-BBSRC COVID rolling fund (BB/V017772/1). GLH was also supported by the BBSRC (BB/T001240/1 and BB/V011278/1), a Royal Society Wolfson Fellowship (RSWF\R1\180013), the NIH (R21AI138074), the EPSRC (EP/V043811/1), the UKRI (20197 and 85336), and the NIHR (NIHR2000907). GLH is affiliated to the National Institute for Health Research Health Protection Research Unit (NIHR HPRU) in Emerging and Zoonotic Infections at University of Liverpool in partnership with Public Health England (PHE), in collaboration with Liverpool School of Tropical Medicine and the University of Oxford. GLH is based at LSTM. The views expressed are those of the author(s) and not necessarily those of the NHS, the NIHR, the Department of Health or Public Health England. USA-WA1/2020 was deposited by the Centre’s for Disease Control and Prevention and obtained through BEI Resources. NIAID, NIH: SARS-Related Coronavirus 2, Isolate, NR-52281. B.1.1.7 (hCoV-19/England/204820464/2020) was obtained through BEI Resources, NIAID, NIH: SARS-Related Coronavirus 2, Isolate hCoV-19/England/204820464/2020, NR-54000, contributed by Bassam Hallis. B.1.351 (hCoV-19/South Africa/KRISP-EC-K005321/2020) was obtained through BEI Resources, NIAID, NIH: SARS-Related Coronavirus 2, Isolate hCoV-19/South Africa/KRISP-EC-K005321/2020, NR-54008, contributed by Alex Sigal and Tulio de Oliveira. P.1 (hCoV-19/Japan/TY7-503/2021) was obtained through BEI Resources, NIAID, NIH: SARS-Related Coronavirus 2, Isolate hCoV-19/Japan/TY7-503/2021 (Brazil P.1), NR-54982, contributed by National Institute of Infectious Diseases.

